# A national crop wild relative checklist for Zimbabwe reveals edible crop wild relative diversity of regional and global importance

**DOI:** 10.1101/2025.07.04.663189

**Authors:** Kudakwashe Mutasa, Onismus Chipfunde, Christopher Chapano, Patience Chatukuta

**Affiliations:** National Herbarium and Botanic Garden, Department of Research and Specialist Services, Harare, Zimbabwe; National Plant Genetic Resources Institute, Department of Research and Specialist Services, Harare, Zimbabwe; Max Planck Institute for Biology Tübingen, Tübingen. Germany

## Abstract

Crop wild relatives (CWRs) are wild plant species genetically related to cultivated crops and represent vital reservoirs of genetic diversity for food security and agricultural resilience. Zimbabwe, with its five floristic regions, hosts substantial plant biodiversity, yet a comprehensive national inventory of its CWRs has been lacking. This study presents a desk-based assessment to develop a national checklist of Zimbabwean CWRs, applying a conceptual framework that evaluates edible CWRs in terms of taxonomic composition, use categories, biogeographic status, extinction risk, breeding potential, and vernacular nomenclature. An integrated approach combining floristic, ecological, and ethnobotanical data was employed to contextualize their utility in food systems and conservation.

Findings reveal that Zimbabwe harbors over 2,700 CWR taxa across more than 100 plant families and nearly 400 genera, related to approximately 260 cultivated crops. Documented uses span food, fodder, medicine, ornamental, and industrial applications, with edible uses comprising nearly 40% of taxa with recorded utility. Edible CWRs span 53 families, predominantly Orchidaceae, Poaceae, Fabaceae, Convolvulaceae, Lamiaceae, and Euphorbiaceae, and serve diverse dietary roles including vegetables, fruits, spices, herbs, cereals, oils, and beverages. Around 90% CWR are native, and one-third are regionally endemic, with the Eastern Highlands region identified as a biodiversity hotspot. Despite their significance, only 30% of edible CWRs have been assessed for extinction risk, and just 0.2% have documented confirmed or potential traits for crop improvement. Vernacular names were recorded in six local languages for 26% of edible taxa, highlighting significant cultural knowledge and integration into local food systems.

This study underscores the underutilized potential of edible CWRs in plant breeding and conservation planning. Their diversity, cultural embeddedness, and adaptive traits present significant opportunities for enhancing food security and advancing sustainable development in Zimbabwe and beyond. Strategic conservation and utilization of these genetic resources are urgently needed.

## Introduction

Crop wild relatives (CWRs)—wild plant species phylogenetically related to cultivated crops— play a critical role in sustaining global food, fodder, medicinal, ornamental, and industrial crop systems (Maxted et al., 2006; Mansfeld and Hanelt, 2001; Raymond, 2006.). A CWR is a wild taxon found (i) in the **gene pool** of a crop where gene exchange is possible or (ii) in the **taxon group** of the crop up to genus level (Maxted et al., 2006). Current discourse around CWRs recognizes them as reservoirs of genetic diversity and adaptive traits vital for enhancing resilience to stress, particularly as agriculture confronts climate change and biodiversity loss (Tanksley and McCouch, 1997). International agreements such as the Cartagena Protocol on Biosafety and the Kunming-Montreal Global Biodiversity Framework highlight the urgency of conserving the genetic diversity within CWR taxa (CBD, 2005; CBD, 2022). The greatest diversity of wild relatives for any agricultural crop is often found in the countries of the crop’s origin and/or domestication, making it the country’s responsibility to conserve this diversity (Maxted et al., 2006; Castañeda-Álvarez et al., 2016).

However, a less emphasized yet equally critical dimension of CWRs is their direct and immediate utility in human nutrition. Many CWRs are actively and routinely consumed as food and provide nutritional benefits to humans and animals alike. In the context of this study, edible CWRs are those CWRs with a documented use as food. Edible CWRs supplement staple diets, buffer communities against seasonal food shortages, and provide dietary diversity (Maroyi, 2011; Chinomona et al., 2024), including as food additives. Additionally, in the rural setting, CWRs support traditional economies to such an extent that their loss would threaten the traditional socio-economy (Makombe, 1994).

Zimbabwe, forms part of the Flora Zambesiaca region and has an estimated 5,930 plant taxa found in a diverse range of vegetation types including Miombo, Mopane, Teak, Acacia, and Terminalia-Combretum woodlands (Mufandaedza, 2002; Mapaura and Timberlake, 2004) which support livelihoods of over 67% of the population (Ministry of Lands, 2019). These ecosystems, in turn, support a high density of CWR taxa, many of which are edible and culturally significant. Yet, despite their ecological and nutritional importance, edible CWRs in Zimbabwe face mounting threats, including habitat loss from land use changes, climate change, commercial agriculture, alien plant species invasions, pest and disease distribution changes, evolving societal attitudes, erosion of traditional knowledge systems, and poor representation in plant collections (Manduna and Vibrans, 2019; Macheka et al., 2022; Magos Brehm et al., 2022).

Traditionally, local communities have developed *in situ* and *ex situ* conservation practices— such as selective foraging, intercropping with cultivated species, and protection during land clearing—that both conserve and propagate edible CWRs (Campbe, 1987; High and Shackleton, 2000; Maroyi, 2011; Macheka et al., 2022). These conservation efforts were mainly done at local level until recently when global attention recognized the threat faced by CWRs in the face of continued deterioration of the environment and effects of climate change. In response to the international agenda of conserving plant genetic resources, the Zimbabwean government, in collaboration with development partners, set out to develop the first comprehensive inventory of CWR recorded in the country, building on the initiative funded through the Southern African Development Community (SADC) Crop Wild Relatives Project. Further support to facilitate digitization of CWRs was obtained from the Max Planck Institute for Biology Tübingen.

This paper presents the first national checklist of crop wild relatives in Zimbabwe. It further analyses the diversity of edible CWRs with regards to their distribution, breeding use, conservation status and traditional importance. This information offers insights into conservation priorities and highlights the importance of Crop Wild Relatives (CWRs) in crop improvement. We emphasize the urgent need to recognize edible CWRs not only as valuable breeding material for the future but also as essential foods for rural diets and livelihoods today. This recognition calls for immediate conservation strategies.

## Methods

The study primarily involved a comprehensive desk review, incorporating qualitative data gathered from a range of sources including herbarium materials, and reputable unpublished and published literature.

### Conceptual framework for the development of the checklist

The national crop wild relatives (CWR) checklist was developed using a structured and systematic approach (Figure 1) adapted from the CWR checklist compilation methodology of Maxted, Brehm, & Kell (2013) (Figure 1).

**Figure 1.**
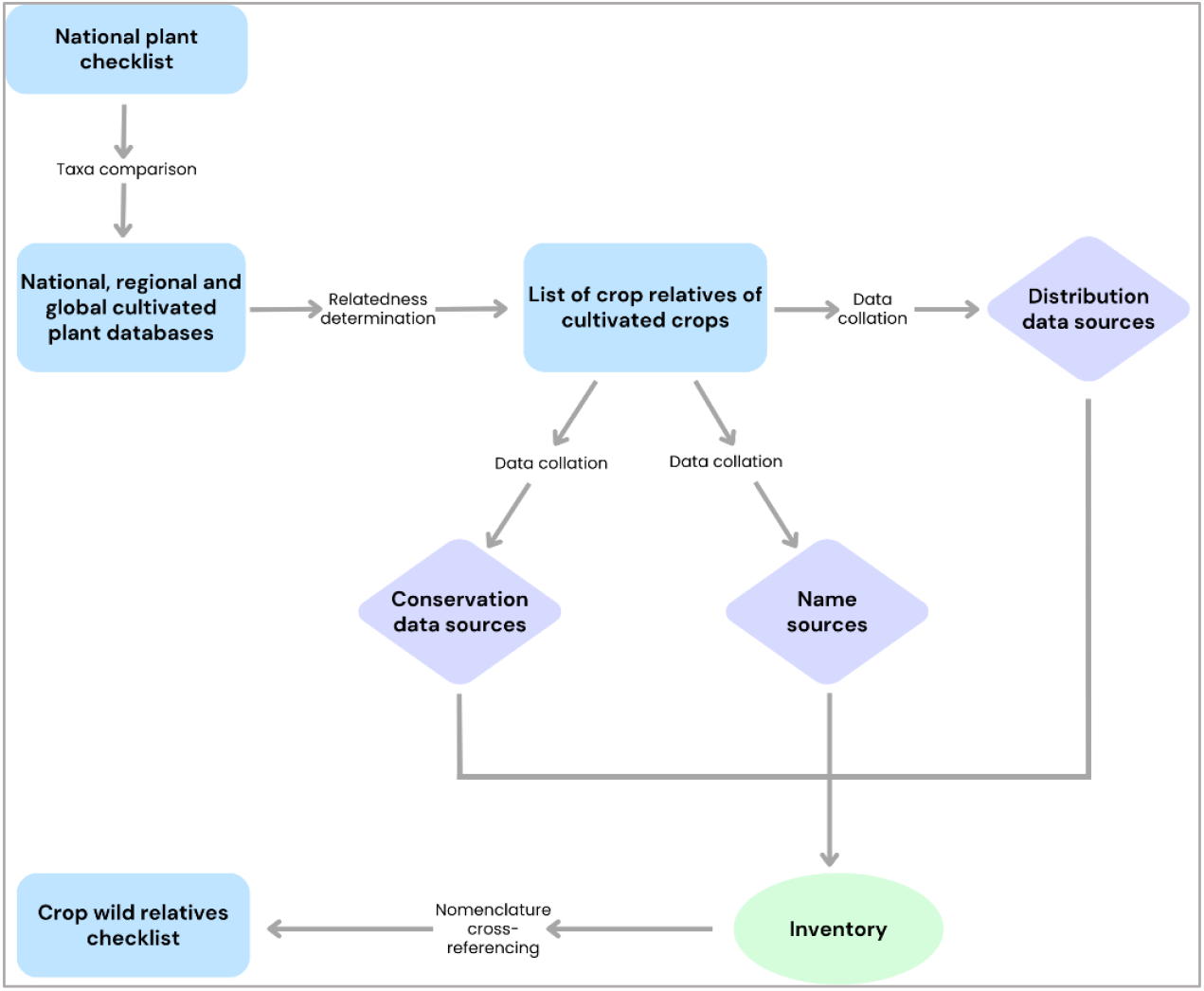
Conceptual framework for development of the Zimbabwe national crop wild relatives checklist.

### Taxonomic Backbone and Initial Dataset Compilation

The foundational taxonomic dataset for the inventory was derived from *A Checklist of Zimbabwean Vascular Plants*, which documents 5,930 vascular plant taxa (Mapaura & Timberlake, 2004). This checklist includes accepted species names that are either indigenous to Zimbabwe or have naturalized within the country’s borders. This primary plant list was cross-referenced against five national, regional, and global databases of cultivated and horticultural crops (Table 1). A taxon in the primary plant list was considered a CWR if its genus matched any of those in the reference databases.

**Table 1.** Databases used for CWR identification through genus-level matching with Zimbabwe’s plant checklist.

|  | Database | Publisher | Year | # of Taxa |
| --- | --- | --- | --- | --- |
| 1 | Mansfeld's World Database of Agricultural and Horticultural Crops | Leibniz Institute of Plant Genetics and Crop Plant Research (IPK) | 2003 | 6,914 cultivated plants excluding forestry and ornamental species |
| 2 | Crop and crop genus list for national CWR checklists and checklist prioritization | S. Kell; SADC CWR project | Unpublished | >1,900 |
| 3 | International Treaty of Plant Genetic Resources for Food and Agriculture – Annex 1 | FAO | 2001 | 131 forage, and major and minor crops |
| 4 | Food and Agriculture Organization Statistics (FAOSTATS) on economic crops | FAOSTAT | 2020 | 101 cultivated plants of economic value |
| 5 | Zimbabwe Genetic Resources and Biotechnology Institute | Ministry of Agriculture, Zimbabwe | Unpublished | 67 cultivated landraces of Zimbabwe |

### Taxonomic Standardization and Synonym Resolution

To ensure consistency in nomenclature, the full list of putative CWR taxa was cross-referenced with the Global Biodiversity Information Facility (GBIF) taxonomic backbone. Accepted species names were retained, and outdated synonyms were updated accordingly. In cases of doubt, plant names were checked for spelling errors or cross-referenced against the International Plant Names Index (IPNI). Taxa with unresolved nomenclature were excluded from the inventory or flagged for further review. In cases where, the revisions in taxa have been done, the genetic relationship with the crop was verified using records from the Harlan and de Wet Crop Wild Relative Inventory and the USDA Germplasm Resources Information Network (USDA, 2025)

### Conservation and Utilization Attributes

For each identified CWR taxon, conservation and utilization information was collated from multiple sources. Endemism data were extracted from the *Endemic Plant Species of Zimbabwe* (Mapaura, 2002), while species threat status at the national level was cross-checked against the Zimbabwe national red data list (Mapaura & Timberlake, 2002). Utilization potential and breeding uses—such as known gene pool associations or crop improvement relevance—were annotated using records from the Harlan and de Wet Crop Wild Relative Inventory and the USDA Germplasm Resources Information Network (USDA, 2025). The gene pool (GP) and taxon group (TG) classification systems (Harlan & de Wet, 1971; Maxted et al., 2006) were applied to define the relationships between wild relatives and their cultivated counterparts.

### Spatial Distribution and Vernacular Knowledge

Species occurrence and distribution data were collated from the national checklist (Mapaura & Timberlake, 2004) for the generalized location and supplemented with geo-referenced records from online biodiversity databases, including the Zimbabwe Flora website (www.zimbabweflora.org; Hyde et al., 2025), GBIF (www.gbif.org), USDA’s GRIN-Global (https://npgsweb.ars-grin.gov/gringlobal), and digital data of national herbarium specimens. These sources enabled mapping of national hotspots of CWR richness.

Common and vernacular plant names were compiled from two national botanical resources (Mullin, 2003; Kwembeya & Takawira, 2002), providing insights into ethnobotanical relevance and community-level knowledge of the species.

## Results

### Composition and Use Categories of Zimbabwe’s Crop Wild Relatives

The comprehensive checklist comprising 2,767 crop wild relative (CWR) taxa in Zimbabwe, spanning native, endemic, and introduced species, represents 123 families, 399 genera, 2,612 species and 2,764 subspecies which are related to 259 cultivated crop species. These taxa represent diverse use categories including food, fodder, medicinal, ornamental, forestry, and industrial applications (Supplementary Information 1). Of these, 1,802 taxa have documented uses, with 997 taxa (36,0%) categorized as edible CWRs (Figure 2).

**Figure 2:**
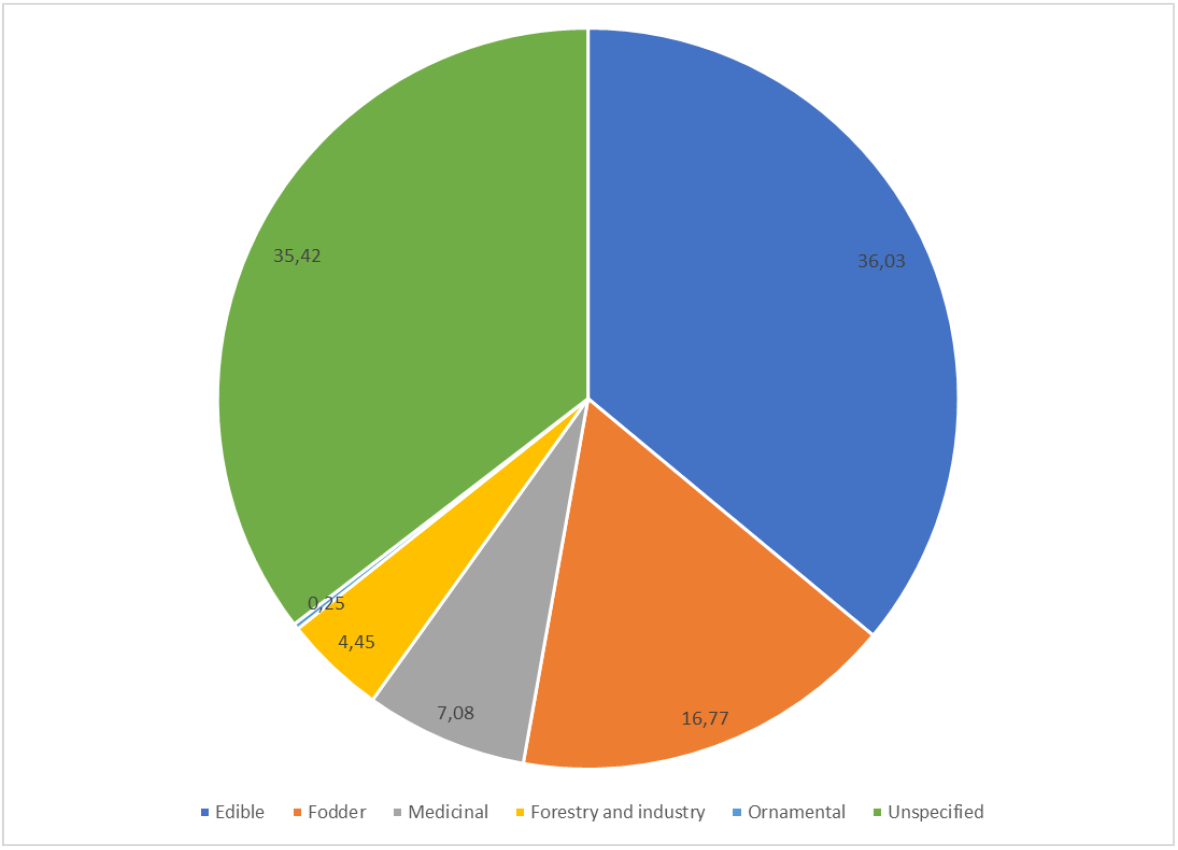
Distribution of CWR by documented use.

### Taxonomic Diversity and Functional Categorization of Edible CWRs

The 997 edible CWRs are distributed across 53 plant families. The most represented families include Orchidaceae (233 taxa), primarily used for flavouring; Poaceae (165 taxa), which encompass cereals and starchy grasses; Fabaceae (105 taxa), key sources of plant-based protein; Convolvulaceae (61 taxa), notably contributing tuber crops; Lamiaceae (48 taxa), known for non-starchy tubers and aromatic and flavouring taxa; and Euphorbiaceae (44 taxa), which provide sweet fruits (Figure 3).

**Figure 3:**
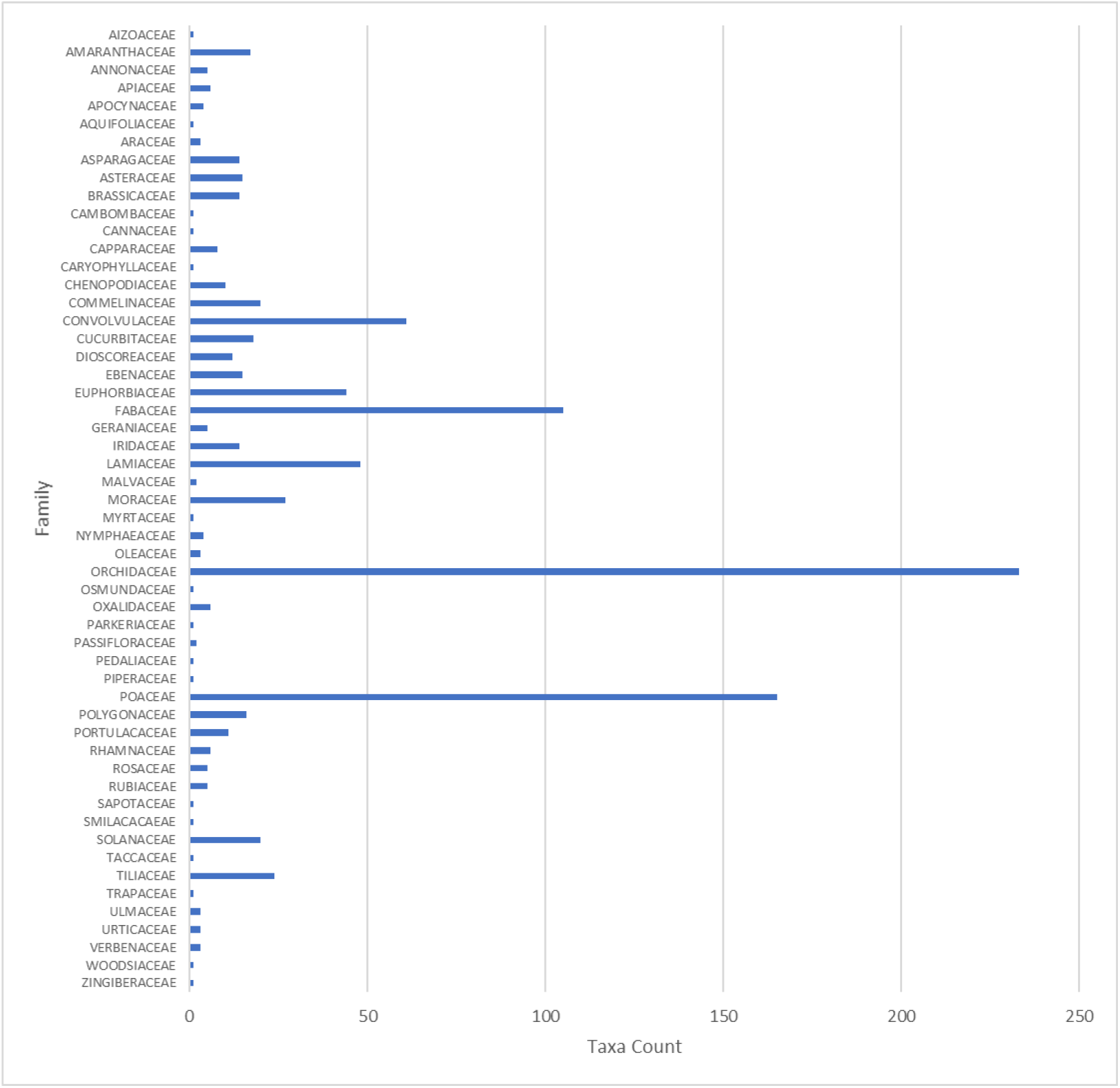
Number of edible CWR taxa per family.

Edible CWRs were further analysed based on their specific food use (Figure 4). Five primary categories—flavouring, vegetables, cereals, tubers, and fruits—account for nearly 80% of edible CWRs. The flavouring category (266 taxa, 26.68%) includes species traditionally used as herbs, spices, and aromatic agents. Tubers (115 taxa, 11.53%) and cereals (158 taxa, 15.84%) reflect the centrality of starch sources in traditional diets. Vegetables (174 taxa, 17.45%) and fruits (82 taxa, 8.22%) are also prominent, underscoring the contribution of CWRs to dietary diversity. Minor but notable use categories include pulses (2.51%), oil crops (2.41%), and beverages (3.71%). Approximately 4% of edible CWRs have no currently specified food use.

**Figure 4:**
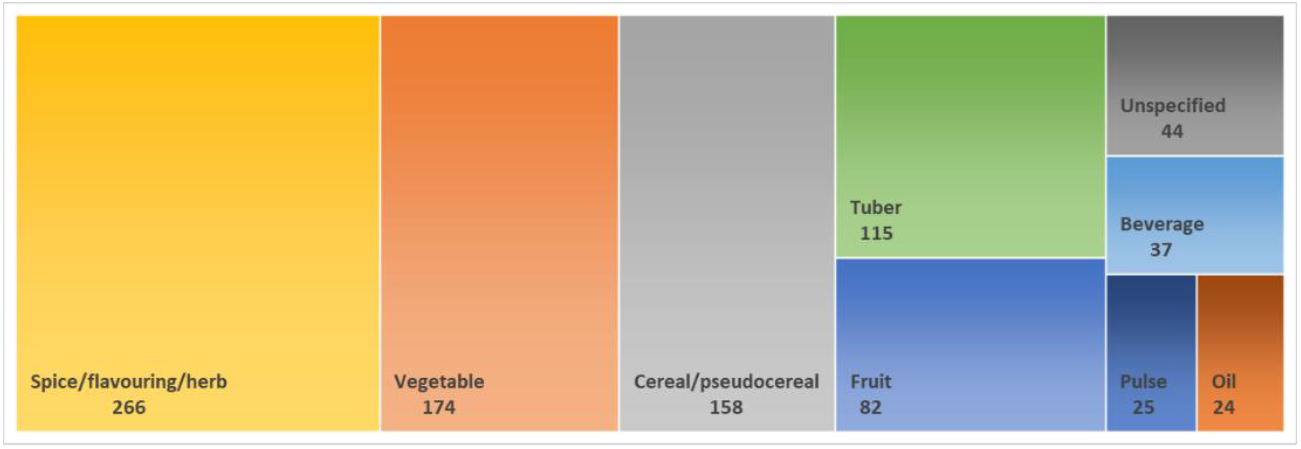
Edible CWR distribution by food use category. Numbers indicate taxa counts in each category.

### Biogeographic Status of Edible CWRs

The country’s rich indigenous diversity in edible CWR is evident with approximately 90% of edible CWRs identified being native to Zimbabwe, while only 9% are exotic or introduced from other countries. Distribution data by floristic region shows that about one-third of taxa in each region are edible taxa (Figure 5). The Eastern highlands presents the region with the highest number of indigenous CWRs while the southern and western regions of the country have the least number.

**Figure 5:**
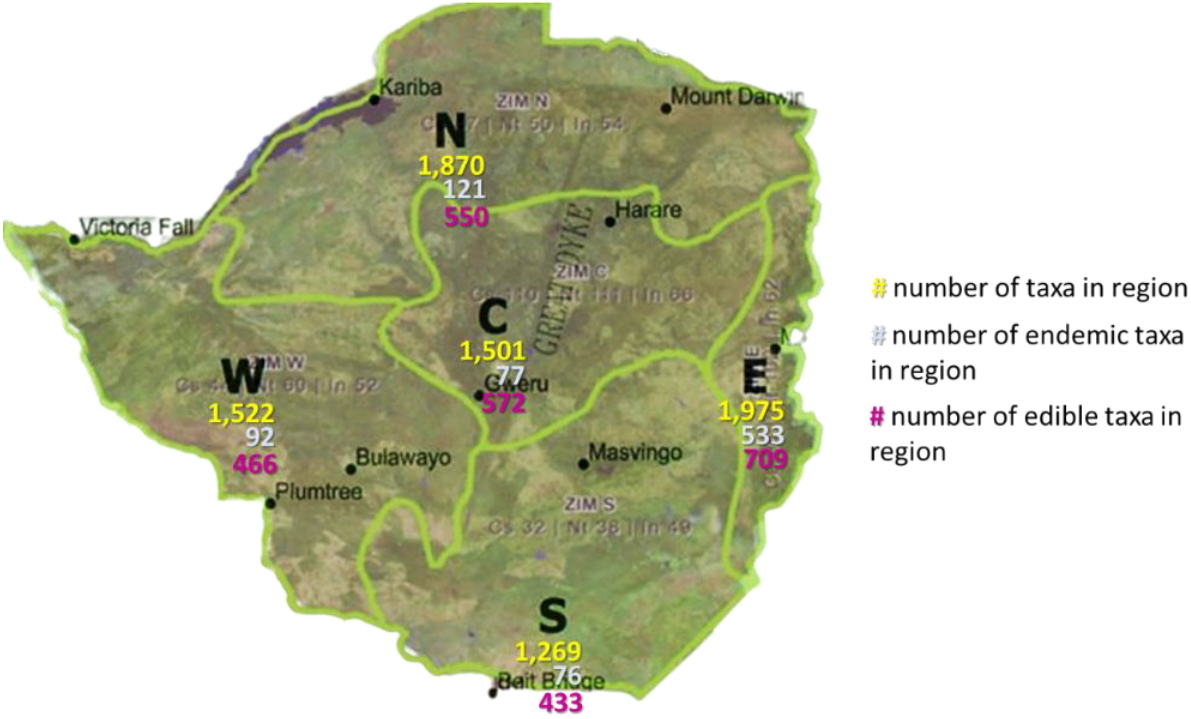
Plant taxa distribution in Zimbabwe by floristic region. Zimbabwe is divided into 5 floristic regions: North (N), West (W), Central (C), East (E) and South (S).

**Figure 6:**
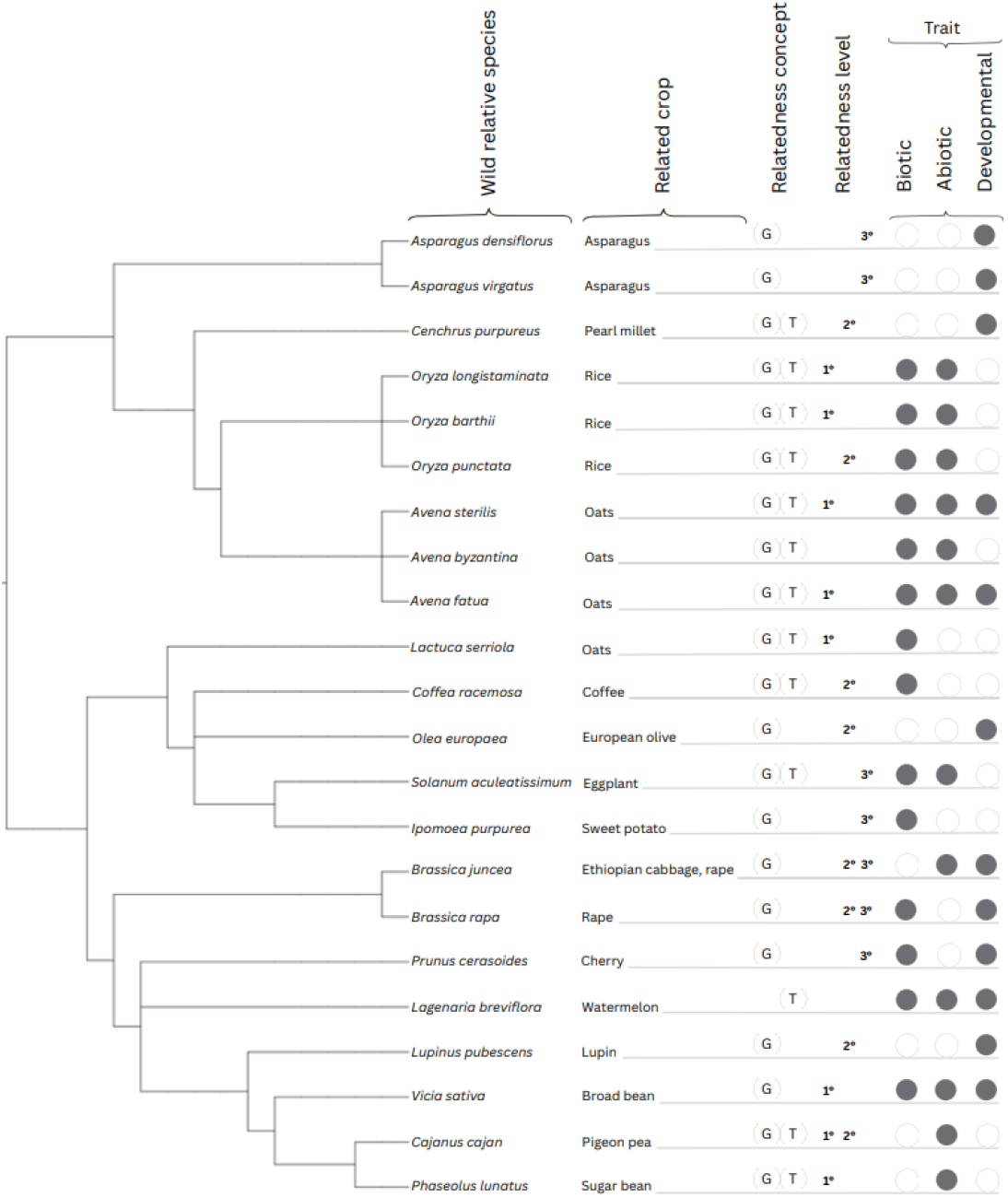
Edible CWR taxa with potential or confirmed useful traits for breeding. G = genepool. T = taxon group. White circle = absence. Grey circle = presence. Biotic = Biotic stress tolerance or resistance. Abiotic = abiotic stress tolerance or resistance. Detailed traits listed in Supplementary information 2. Phylogenetic tree visualized in iTOL v6 (Letunic and Bork, 2024).

**Figure 7:**
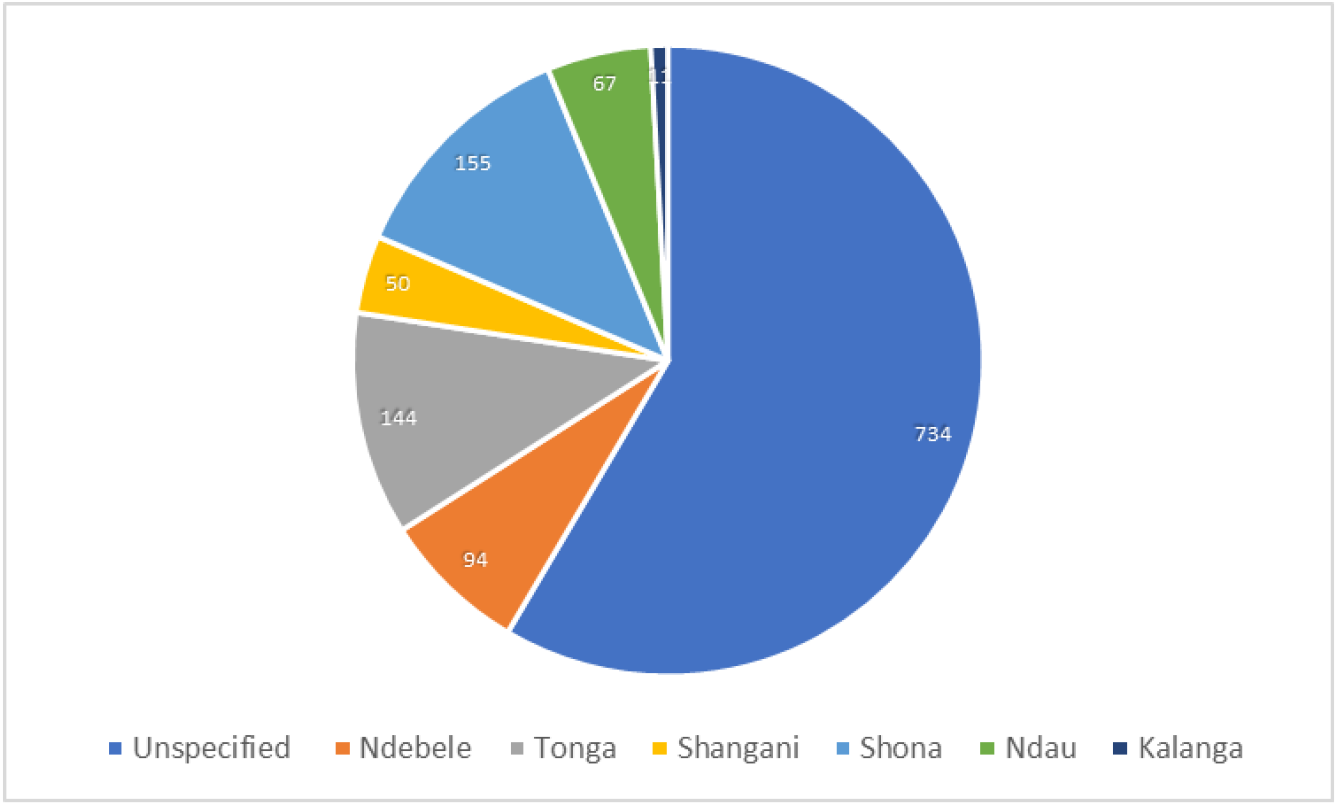
Distribution of vernacular languages of Zimbabwe among edible CWR taxa. Some taxa have multiple names in different languages and dialects.

Relative high endemism was recorded. Endemic plants are plants with a restricted geographic region or are known to be found in a specific locality. The eastern region has 533 endemic taxa, and also has the highest number of total taxa and the most edible taxa. The remaining 4 regions each have less than 125 endemic taxa. The high species richness, endemism of over 4 times higher than any other region, and high risk of extinction in the Eastern Highlands of Zimbabwe highlights this area as a critical biodiversity hotspot due to their unique and at-risk flora. The north, west, central and southern regions support significantly diverse vegetation and therefore require region-specific conservation strategies.

### Conservation Assessment and Threat Status

Conservation assessments for edible CWRs are outdated and limited. Of the 997 edible CWR taxa, only 335 taxa (33.6%) had threat assessments conducted in 2002—14 at the national level and 321 at the regional level (IUCN Red Data List, 2025). A total of 662 taxa (66.4%) lack any known threat status assessment, reflecting a major knowledge gap. Only 16 edible CWRs are currently listed in the IUCN Red List of Threatened Species, within the categories *Least Concern, Vulnerable, Near-threatened, Endangered* and *Critically Endangered* (Table 2). No edible CWRs were classified as extinct or extinct in the wild.

**Table 2:** Edible CWR taxa with known extinction risk status based on 2002 assessments.

| Status | Family | Genus | Species |
| --- | --- | --- | --- |
| **Least concern** | LAMIACEAE | Plectranthus | Porphyranthus |
| **Vulnerable** | MORACEAE | Ficus | Scassellatii |
|  | RUBIACEAE | Coffea | Ligustroides |
|  | RUBIACEAE | Coffea | Mufindiensis |
|  | CONVOLVULACEAE | Ipomoea | Verrucisepala |
| **Near threatened** | LAMIACEAE | Plectranthus | Caudatus |
|  | MORACEAE | Ficus | exasperate |
|  | POACEAE | Eragrostis | desolata |
| **Endangered** | RUBIACEAE | Coffea | ligustroides |
|  | RUBIACEAE | Coffea | zanguebariae |
|  | CONVOLVULACEAE | Ipomoea | verrucisepala |
|  | MORACEAE | Ficus | vallis-choudae |
|  | ORCHIDACEAE | Vanilla | polylepis |
| **Critically endangered** | MORACEAE | Ficus | bubu |
|  | MORACEAE | Ficus | fischeri |
|  | MORACEAE | Ficus | modesta |

### Breeding Potential of Edible CWRs

A total of **23 edible CWR taxa** were identified as having **confirmed or potential traits useful for crop improvement** (Table 3). These traits include:

- **Biotic stress resistance** (e.g., resistance to fungal diseases, bacterial blight, insect pests).
- **Abiotic stress tolerance** (e.g., drought, frost, cold, heat, soil salinity).
- **Agronomic traits** (e.g., seed weight, yield, panicle length, male sterility for hybrid breeding).

Several of these taxa (e.g., *Oryza barthii, Avena fatua, Cajanus cajan*) are part of **primary or secondary gene pools** for major crops, suggesting high utility for breeding programs. The data underscore Zimbabwe’s potential for contribution to global food security through genetic diversity in CWRs.

### Vernacular Knowledge and Cultural Relevance

Vernacular names were documented for **263 edible CWR taxa**, while **46% of the total taxa have known common names in English**. About 74% of edible CWRs have no documented vernacular names. Vernacular names span major local languages with 155 edible CW taxa having chiShona names, 94 isiNdebele, 67 chiNdau, 50 chiShangani, 11 tjiKalanga, and 144 chiTonga. Among the chiShona names, various dialects (chiZezuru, chiKaranga, chiManyika and chiZezuru) have been recorded. Many taxa have vernacular names in more than one language and a low number of taxa have vernacular names in only 1 language. Vernacular naming patterns can reveal local use, cultural importance, and regional familiarity. They may also point to regional adaptations and the high value placed on these taxa for nutrition.

## Discussion

This study presents a comprehensive checklist of crop wild relatives (CWRs) in Zimbabwe, documenting 2,767 taxa, of which 997 are classified as edible. Using an integrated approach— linking floristic, ecological, and ethnobotanical data—we generated a national checklist that not only catalogs Zimbabwe’s CWRs but also contextualizes their utility in local food systems and their conservation value. This checklist represents a foundational tool for national plant genetic resource management, research prioritization, and policy development.

When compared with other countries in the Southern African Development Community (SADC) region, Zimbabwe has one of the most diverse CWR inventories. SADC documents approximately 1,900 CWR species related to 64 priority food and beverage crops (Allen et al., 2019), while Malawi, Mauritius, Zambia, and South Africa report 446, 528, 459, and 1,859 taxa respectively (Mponya et al., 2020; Bissesur et al., 2019; Ng’uni et al., 2017; Holness et al., 2019). Zimbabwe’s comparatively higher figure reflects both its floristic diversity, and the depth of national plant assessments and ethnobotanical knowledge. The proportion of introduced species reflects the history of plant exchanges during migration, trade and colonization (National Research Council, 2002; Hulme, 2021).

The checklist reveals the multifunctional values of these plants, as species for food consumption or for contributing traits to food crops, as livestock feed, as traditional healthcare, as industrial raw materials, and as tools for aesthetic uses. Almost two-thirds of the CWR taxa have documented uses, indicating the local population’s high level of ethnobotanical knowledge and ability to integrate these plants into their livelihoods. This utilization rate suggests that local knowledge systems actively engage with and benefit from CWRs, reinforcing the utility and embeddedness of CWRs (Mujuru et al., 2020; de Medeiros et al., 2021). Given that the uses were documented at global level, this reflects on the potential of Zimbabwe’s local genetic resources for contributing to global food security and economic benefit.

Over half of the usable CWRs are edible plants of importance in local diets and are therefore tied to food security and have potential for diversification of agricultural production. Functional classification of edible CWRs provides an understanding that CWRs are deeply embedded in traditional culture for nutrient intake and for low-input farming systems (IPES-Food, 2024). Local cereals and tubers tend to be climate-resilient and can therefore support climate adaptation efforts (Zenda et al., 2021; Glatzel et al., 2025). Vegetables, fruits and flavor-related crops provide dietary diversity, micronutrients and opportunity for income generation. While few in number, pulses and oil crops provide essential fats and proteins, whereas beverage-related crops hold cultural and commercial value as ingredients in traditional drinks (Glatzel et al., 2025). The presence of taxa with unspecified uses suggests absence of traditional use or lack of documentation of such uses, thus highlighting an opportunity for further research in underexplored regions or communities.

In addition, the diversity of vernacular names across six major local languages reflects both regional use-patterns and the linguistic richness of the communities involved. About 26% of edible CWRs possess vernacular names, with the majority having vernacular names in multiple languages. The majority of the edible CWRs lack recorded vernacular names, suggesting either poor documentation or low levels of familiarity and utilization locally. Relatively few (18%) are restricted to a single language, implying some degree of cross-cultural utilization and shared ethnobotanical knowledge. However, taxa with vernacular names confined to one language may represent species with restricted geographic distributions and localized uses. Vernacular naming patterns provide insights into ecological familiarity, historical use and regional adaptations of edible CWRs. A compelling case is finger millet (*Eleusine coracana*) which boasts 32 names in chiShona dialects alone; these names distinguish finger millet varieties according to their phenotypic characteristics such as head colour, maturing time or grain size (Mullin, 2003), indicating an intimate knowledge of plant varieties by the local people and highlighting the importance of vernacular languages in preserving and transmitting agronomic knowledge that may be absent or under-represented in global databases. The close alignment of food use and vernacular names with phenotypic traits targeted in crop breeding allows for the convergence of traditional knowledge and scientific trait characterization, offering pathways for community-based conservation and participatory breeding.

The distribution of the edible CWRs reported in this paper across 53 plant families signifies broad taxonomic diversity and suggests ecological adaptation across varied environments. These CWRs may provide a large pool of traits for crop improvement. About 7,040 edible plants are recognized for their potential as future foods (Antonelli et al., 2023), although only 15 plants provide 90% of global food intake by humans. Globally, the most species-rich families with respect to edible species are the Fabaceae, Arecaceae, Poaceae, Malvaceae and Asteraceae; and with respect to major food crop species, Rosaceae, Fabaceae, Dioscoreaceae, Poaceae and Malvaceae (Ulian et al., 2020). Surprisingly, the most represented family is Orchidaceae, which highlights the importance of non-staple uses in CWR plant use. The Orchidaceae CWR are potentially an under-researched economic resource for food flavouring. The grass family (Poaceae) CWR are well-adapted to marginal environments, and are therefore critical for provision of starch under climate change. The most economically important dicotyledon family (Bisby, 1994), the pulses (Fabaceae), produces protein-rich seeds and can fix nitrogen in the soil, and is therefore beneficial for protein intake and improved soil fertility in low-input, climate-resilient farming (McKey, 1994; Bennet, 2003; Koenen et al., 2019).

Convolvulaceae include wild relatives of potato and other tubers which are suited for poor soils and water-limited areas (Simões et al., 2024). Euphorbiaceae includes species with sweet or fleshy fruits, and also toxic species (Hohmann and Molnár, 2004; Liu et al., 2024), highlighting traditional knowledge in nutritional value, and safe preparation and use. The Poaceae and Fabaceae families are closely related to global food crops such as rice, wheat, maize, soya and common bean, making them top candidates for pre-breeding programs. Under-researched families such as the Orchidaceae and Lamiaceae may hold untapped economic potential in niche markets such as spices and organic products.

Edible CWRs are broadly accessible across Zimbabwe, making up about one-third of taxa in each floristic region. The implications of the dominance of native edible CWR species are that local ecological adaptation facilitates *in situ* conservation (Berkes et al., 2000; Meek et al., 2022), historical cultural integration in traditional diets and practices (Wootton and Lyver, 2024), and crop improvement using locally-adapted traits. Only 9% of edible CWRs are non-native and they require careful management to contribute new traits and uses, or to prevent ecological competition or invasiveness.

The Eastern Highlands region stands out as a critical biodiversity hotspot due to its species richness, high endemism and highest number of edible CWR taxa. The region also has elevated risk of extinction due to habitat loss, climate change and other threats. The threats include natural disasters, such as Cyclone Idai which, in 2019, devastated natural forests and plantations in the Eastern Highlands (Chikodzi and Tembani, 2021). The Eastern Highlands should therefore be a national conservation priority for *in situ* CWR conservation, protected area expansion, and research. Although not as rich in endemics as the Eastern Highlands, the north, west, central and south floristic regions support diverse vegetation and high CWR representation, and demand localized region-specific interventions that are unique to their flora and threats. The west and south regions are particularly vulnerable to drought conditions and rely more on edible CWRs that the rest of the country due to their poor food security status.

The finding that only about 34% of edible CWRs have been assessed for threat status as far back as 2002 reveals a critical and major gap in current, evidence-based conservation planning because a large proportion of edible CWRs are invisible to conservation decision-making. Additionally, the low representation of edible CWRs on the IUCN Red List suggests that economically dominant crops are preferentially researched over regionally-important species and that there are systemic biases resulting in underrepresentation of African flora in global conservation assessments (Meyer et al., 2016). The lack of assessments may reflect insufficient funding and poor data infrastructure for plant conservation in Zimbabwe. There are no recorded extinctions or near-extinctions yet, but this may be due to inadequate monitoring. Given extensive land use changes, forest loss, and natural disasters since 2002, these numbers may underrepresent the current extinction risk, emphasizing the urgent need for a national reassessment to update the conservation status using current IUCN criteria and to identify priority taxa for conservation action. Many unassessed species may be growing in isolated or under-threat habitats, and may require community-based conservation in addition to seed banking and *ex situ* living collections. Collaborative partnerships with herbaria, botanic gardens, universities, research institutes and local communities can help bridge this gap. Strategic investment in research, reassessment and conservation infrastructure is essential to safeguard these invaluable plant genetic resources.

Only a small subset (∼2.3%) of edible CWRs has confirmed or potential traits for crop improvement and these taxa hold disproportionately high value for contribution to biotic and abiotic stress tolerance and resistance, as well as agronomic improvements. These traits are essential for climate-resilient crops that can withstand emerging challenges in both subsistence and commercial agriculture. The presence of these high-value breeding traits provides Zimbabwe with a unique opportunity to position itself as a centre for CWR conservation and pre-breeding, leveraging multi-omics and breeding to unlock their full potential. Furthermore, the small size of the subset reflects poor data availability rather than the true extent of breeding potential, and therefore there is a need to systematically evaluate more edible CWRs for beneficial traits.

The ongoing digitisation effort at the National Herbarium of Zimbabwe shows that the government holds a substantial number of historical specimens representing 2,497 taxa. The national gene bank conserves approximately 6,500 accessions of cultivated plant genetic resources including a very small number of accessions of CWRs (less than 1%) for Oryza, Sorghum and Cucumis species. Critically, the absence of most CWRs from the national gene bank inhibits their incorporation into national food security initiatives. Although 23 CWR taxa with useful breeding traits have been identified in this study, the actual number of taxa historically used for crop improvement in Zimbabwe is not known, suggesting under-documentation and underutilization. The importance of conserving and utilizing these CWRs becomes evident when placed in the context of Zimbabwe’s agricultural economy, which contributes up to 15% to the national GDP and employs 70% of the labour force (Ministry of Agriculture, 2019). Major crop groups cultivated in the country include cereals, legumes, fruits, tobacco, and forestry species, many of which have CWRs that are documented in this study and could be useful for development of climate-resilient crop varieties. Importantly, the presence of use-records for such a high proportion of CWRs supports the need to integrate indigenous knowledge systems into formal conservation and agricultural planning. The existing national policy framework which includes the ten-year National Strategy and Action Plan for Plant Genetic Resources for Food and Agriculture produced by the Ministry responsible for agriculture in 2022 recognises CWRs as a strategic priority group of plant species for management of plant genetic resources.

An important but unacknowledged threat to edible CWRs is the erosion of traditional knowledge and increasing preference for commercial hybrid crops. Negative perceptions of CWRs from colonial times have persisted to date and have led to a decline in the use and value of wild relatives. Edible CWRs are commonly foraged in rural areas and are often associated with poverty and food insecurity, especially during times of hardship (Manduna and Vibrans, 2019). Poor transport infrastructure restricts the distribution of edible CWRs to their immediate catchment areas, leading to low awareness of these taxa outside of local communities (Maroyi, 2011). Moreover, academic perspectives have sometimes categorized these valuable plants as weeds within monocultural farming systems (Mavengahama, 2013). These negative cultural and scientific perceptions limit the visibility and valuation of CWRs in both policy and practice.

However, the negative perception is slowly changing as growing global trends towards healthier and more organic diets have seen gentrification of certain edible CWRs. Plants once considered “poor people’s crops,” such as *Fadogia ancylantha* (makoni tea), *Myrothamnus flabellifolius* (resurrection bush), *Eleusine coracana* (finger millet), and *Pennisetum glaucum* (pearl millet), are now being marketed as premium superfoods. Similarly, indigenous fruit tree species such as baobab and marula are gaining prominence in nature-based industries. Medicinal uses of CWRs are also gaining renewed attention, especially in the face of rising healthcare costs and limited access to conventional medicine. However, these changes bring risks—chiefly, ecological degradation through overharvesting, exclusion of local communities from benefits, and market-driven commodification that may reduce equitable access. This underscores the need for better documentation and integration of ethnobotanical knowledge into national health strategies.

Public awareness campaigns are essential to reversing negative perceptions and encouraging conservation. These should be targeted across a wide range of stakeholders, including research institutions, agricultural extension workers, national park staff, policymakers, and local communities. Notably, gender-sensitive approaches are critical. Research indicates that women, due to their roles in food preparation and household provisioning, hold more extensive knowledge about leafy vegetables, indigenous fruits, and edible roots (Macheka et al., 2022). Thus, ensuring gender inclusivity in knowledge documentation initiatives is key to capturing a comprehensive understanding of CWR usage.

Finally, the findings of this study point to immense potential for edible CWRs to contribute meaningfully to national and regional food systems, particularly in an era of climate uncertainty. The unavailability of recent information on plant diversity assessments in the face of ecological disturbance due to human activity points to high risk of extinction of CWRs. Therefore, national plant surveys, inventories, conservation, utilization, and equitable access must be prioritized to ensure that the benefits of these invaluable resources are not lost but instead leveraged for future food security and sustainable development.

